# High Throughput Intact Protein Analysis for Drug Discovery Using Infrared Matrix-Assisted Laser Desorption Electrospray Ionization Mass Spectrometry

**DOI:** 10.1101/2021.11.08.467755

**Authors:** Fan Pu, Scott A. Ugrin, Andrew J. Radosevich, David Chang-Yen, James W. Sawicki, Nari N. Talaty, Nathaniel L. Elsen, Jon D. Williams

## Abstract

Mass spectrometry (MS) is the primary analytical tool used to characterize proteins within the biopharmaceutical industry. Electrospray ionization (ESI) coupled to liquid chromatography (LC) is the current gold standard for intact protein analysis. However, inherent speed limitations of LC/MS prevent analysis of large sample numbers (>1000) in a day. Infrared matrix-assisted laser desorption electrospray ionization (IR-MALDESI-MS), an ambient ionization MS technology, has recently been established as a platform for high throughput small molecule analysis. Here we report the applications of such a system for the analysis of intact proteins commonly performed within the drug discovery process. A wide molecular weight range of proteins 10 – 150 kDa was detected on the system with improved tolerance to salts and buffers compared to ESI. With high concentrations and model proteins, a sample rate up to 22 Hz was obtained. For proteins at low concentrations and in buffers used in commonly employed assays, robust data at a sample rate of 1.5 Hz was achieved, which is ∼ 22x faster than current technologies used for high throughput ESI-MS-based protein assays. In addition, two multiplexed plate-based high throughput sample cleanup methods were coupled to IR-MALDESI-MS to enable analysis of samples containing excessive amounts of salts and buffers without fully compromising productivity. Example experiments, which leverage the speed of the IR-MALDESI-MS system to monitor NISTmAb reduction, protein autophosphorylation and compound binding kinetics in near real-time, are demonstrated.

**Graphical Abstract:** 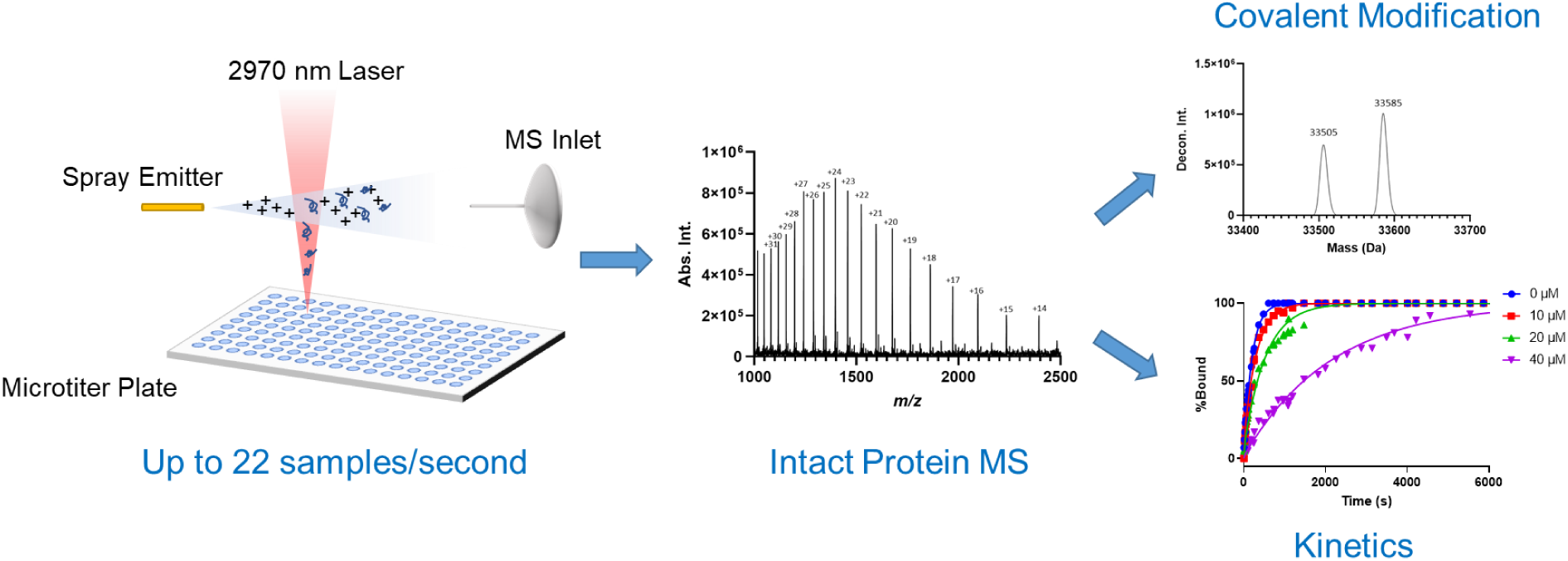

## Introduction

A wide variety of recombinant proteins are produced for drug discovery in the biopharmaceutical industry that require identity confirmation and characterization. Target proteins are developed as reagents for small molecule drug discovery to perform screening assays to find hit and lead compounds and to conduct structural biology studies for structure guided design. The amount and extent of target protein modification by phosphorylation and other post-translational modifications or covalent small molecule inhibitors are determined to ensure protein is in an active form or to enable finding new covalent hit compounds. Therapeutic proteins, such as monoclonal antibodies, dual variable domain antibodies, bispecific antibodies, and fusion proteins, are typically glycosylated during expression. The type and relative amount of glycosylation, along with confirmation of protein identity, need to be determined to ensure accurate and consistent production of protein forms. Many of these therapeutic proteins can then be conjugated with drug and a linker to form an antibody drug conjugate (ADC). The extent of conjugation, given as the drug-to-antibody ratio (DAR) is typically measured by mass spectrometry (MS).

The prevalence of MS in the characterization of both target and therapeutic proteins can find its roots in the introduction of electrospray ionization (ESI)^1^ and matrix assisted laser desorption ionization (MALDI)^2^ techniques, which still remain the primary methods for ionizing proteins.^3^ Vacuum source MALDI can be used to analyze proteins rapidly at a sample rate of >1 Hz (i.e., samples per second) when coupled to a linear time-of-flight (TOF) mass spectrometer and produces primarily singly charged ions, which enables direct determination of molecular weight. However, proteins are typically desalted and crystallized with matrix before ionization, which limits reaction monitoring from being conducted in real-time. These instruments have low mass resolution for high mass singly charged ions, making them insufficient to resolve small mass modifications on large proteins. In contrast, ESI produces multiply charged ions, requiring the use of a deconvolution algorithm to transform the signal from the *m/z* to the mass domain. Despite this, proteins produced by ESI can be coupled to higher resolution mass spectrometers such as orthogonal acceleration TOF (oaTOF), orbitrap (OT), or Fourier Transform – Ion Cyclotron Resonance (FT-ICR) instruments which produce sufficient resolution to detect small mass changes on large proteins.

ESI sources are typically coupled to reversed-phase liquid chromatography (LC) to enable on-line desalting and, if necessary, temporal separation of protein species. For simple recombinant proteins that are present in their native salt / buffer conditions, LC-MS runs of 1 - 5 min can be conducted routinely using <10 pmol of material injected onto the column and be set-up as a part of a fully integrated protein open-access LC-MS system.^4^ These systems are robust and useful for the analysis of up to 384 samples. However, the analysis of 1000 or more samples per day requires the use of a faster sample delivery coupled with a MS-compatible ionization technology.

Progress toward high-throughput protein MS analysis has been achieved by coupling alternative sample introduction devices to an ESI source. Campuzano et al., demonstrated using a RapidFire HT-MS system coupled to an oaTOF system to achieve a sampling rate of 0.05 Hz^5^. They utilized this system to screen the reactivity of 1000 acrylamide compounds after incubation with a 19.5 kDa protein. Sawyer et al., developed a high throughput antibody screening platform with a RapidFire system coupled to an oaTOF, which had a 0.04 Hz sample (injection to injection) rate and was capable of running ∼3400 samples per day.^6^ They also showed that coupling of the RapidFire to an Orbitrap Exploris 480 could be done and led to better resolution of protein glycoforms. Another sample introduction device, the SampleStream, has been developed, which can deliver denatured or native state proteins to a MS system at 0.067 Hz.^7^

For most biochemical and cellular screening assays, sufficient signal-to-noise can be achieved by techniques that do not require a separation step prior to introduction to the mass spectrometer and can analyze at sample rates of 1 – 5 Hz. Some of these techniques include: acoustic mist ionization^8^, acoustic droplet ejection– open port interface^9^, desorption electrospray ionization (DESI)^10^, liquid atmospheric pressure matrix-assisted laser desorption/ionization (LAP-MALDI)^11,12^, and infrared matrix-assisted laser desorption electrospray ionization (IR-MALDESI).^13^ Recently, extremely fast sample rates up to 60 Hz have been demonstrated by LAP-MALDI when interfaced to a Q-TOF system^14^ and 22.7 Hz for IR-MALDESI connected to an OT hybrid instrument.^15^

Many studies have been conducted that demonstrate the utility of ambient ionization techniques in the analysis proteins.^16-28^ DESI has been demonstrated to analyze denatured proteins from 12 kDa to 150 kDa.^16-18^ Specifically, an experiment was conducted to analyze proteins directly from tissue sections.^19^ Native MS with DESI has also been reported recently.^20-21^ With nano-DESI, MS imaging of proteins in tissue has also been demonstrated.^22-23^ Cooper et al. reported the first native-MS imaging of proteins using nanoDESI.^24^ Peng et al. demonstrated with electrospray-assisted laser desorption/ionization (ELDI)^25^ the ability to ionize proteins up to 80 kDa. Vertes et al., using LAESI (laser ablation electrospray ionization), were able to analyze bovine serum albumin (BSA)^26^ and early work by Muddiman et al. demonstrated that MALDESI was capable of intact and top-down protein analysis.^27-28^

Motivated by a need for higher throughput mass spectrometry-based protein assays used in drug discovery, we set out to demonstrate the performance of our recently constructed high-throughput IR-MALDESI source in protein characterization assays. The source described here uses the same geometry, features, and components of the mass spectrometry imaging IR-MALDESI source^29^, but is engineered for direct sampling from industry standard 96-, 384-, or 1536-well plates.^13^ We present in this paper the system performance of our IR-MALDESI-MS by assessing the range of proteins that can be analyzed, detection limits and sampling rate achieved at various resolving power settings, along with studies showing how matrix components affect the signal-to-noise. Additionally, plate-based methods to remove interferents and increase signal-to-noise are demonstrated. Finally, we demonstrate the application of the system in the performance of useful protein-centric kinetics measurements that are essential to the drug discovery process -autophosphorylation, slow and fast compound binding kinetics, and covalent compound additions -that are made possible by the rapid analysis speed of IR-MALDESI-MS.

## Methods and Materials

### Materials

All chemicals except the following were purchased from Sigma-Aldrich: ultrapure ATP was purchased from Promega, 1 M HEPES pH 8 was purchased from Teknova. The following protein standards were purchased from Sigma-Aldrich: cytochrome C (C2037, from bovine heart), carbonic anhydrase (C3934, from bovine erythrocytes), recombinant human serum albumin (rHSA, A6608) and myoglobin (M0630, from equine skeletal muscle).

Drug compounds were obtained from AbbVie compound repository. C-18 BcMag magnetic beads were purchased from Bioclone, C-4 Top-Tip™ and C-4 Lab-in-a-Plate™ desalting devices were purchased from Glygen. Pierce Protein A magnetic beads, Zeba Spin Column (7K MWCO) and protein concentrator (10K MWCO) were purchased from ThermoFisher. BTK kinase domain (387-659) and HCV-La protein were obtained from AbbVie. BTK protein was concentrated to ca. 100 uM and buffer exchanged to 5 mM HEPES pH 7.5 using protein concentrator. La protein used for desalting studies was ^13^C labeled protein produced at AbbVie in a buffer consisting of 25 mM Tris pH 7.5 and 150 mM NaCl. All protein analysis was performed on 384-well small volume HiBase microplates (Greiner Bio-One, 784075) except for desalting experiments. Protein desalting on Blue®Washer (BlueCatBio) and elution from slit plate were performed using 384 well Hard-Shell PCR plates (Bio-Rad, HSP-3805), the PCR plates were used for IR-MALDESI-MS readout directly without transfer.

### Instrumentation

The high throughput IR-MALDESI-MS system was described previously^13^. Briefly, a 2970 nm laser was used to desorb neutral analytes from samples and an orthogonal electrospray emitter aligned to the MS inlet was used to ionize the resulting neutral plume. The IR-MALDESI source was coupled to a QExactive HF-X (ThermoFisher) mass spectrometer. An extended ion transfer tube with custom cartridge heater and motorized stage were used to accommodate analysis of standard microtiter plates.

### MS Conditions

A resolving power (RP) of 7, 500 (FWHM at *m/z* = 200) was adopted for all assays conducted unless otherwise stated to achieve maximum sensitivity with *m/z* ranges selected based on the protein of interest. A spray voltage of 4 kV was supplied to the ESI solvent, which was 80/20 methanol/water v/v with 0.1% formic acid. The ESI flow rate was 2 µL/min All analyses were performed in positive ion mode with Automatic Gain Control (AGC) turned off. The ion transfer tube temperature was set at 400 °C and the temperature of the ion transfer tube extension was set at 120 °C.

### BTK Experiments

The BTK phosphorylation experiment was performed in a buffer composed of 5 mM HEPES pH 7.5, 0.5 mM MgCl_2_, 75 mM ammonium acetate and 0.5 mM ATP with 50 µM of BTK was used.

The BTK binding experiments were performed in 5mM HEPES pH 7.5. 10 mM compounds stock solutions were diluted to desired concentrations with the buffer. 10 µL of 35 µM BTK was first added to wells then an equal volume of 2X compound solution was added to initiate reaction. For denaturing BTK, an equal volume of 1% formic acid was added to the BTK-compound solution.

IR-MALDESI-MS readout was performed from the buffer directly and in real-time, with a time resolution of 3 s.

### Protein Desalting with BcMag™ Magnetic Beads

BCMag™ C18 magnetic beads (Bioclone) were suspended into 50/50 methanol/water v/v at 50 mg/mL and stored at 4 °C. 10 µL/well of the stock mixture was added to plates and storage solution was evacuated on the Blue®Washer (BlueCatBio) on a magnetic carrier. 30 µL/well equilibration buffer (0.5% trifluoroacetic acid, 95/5 water/acetonitrile v/v) was added and removed to equilibrate magbeads. 30 µL protein sample was mixed with 10 µL sampling binding buffer (2% trifluoroacetic acid, 95/5 water/acetonitrile v/v) and incubated for 1 min before adding to working plate. Washing was performed three times with equilibration buffer followed by 30 µL of elution buffer (50/50 acetonitrile/water) to release the protein. Desalted samples were eluted to a PCR plate for IR-MALDESI-MS analyses.

### Protein Desalting with Lab-in-a-Plate™

Lab-in-a-Plate™ (Glygen), is a 384-well plate for protein desalting, based on the Top-Tip™ technology, was packed with C4 stationary plate – P/N THSC04), was procured. The plate was first equilibrated with elution buffer (0.1% formic acid, 50/50 acetonitrile/water) for three times followed by three equilibration cycles with binding buffer (0.1% formic acid). Protein samples were added after equilibration and spun down, followed by three washes with binding buffer and elute with elution buffer. Each wash/equilibration cycle involved centrifuge at 1000g for 1 min except for the last sample elution step, where the plate was centrifuged at 1000g for 2 min.

### Data Analysis

Low resolving power protein MS deconvolution was performed using ReSpect algorithm in Biopharma Finder (ThermoFisher), Xtract was used for resolving power > 60, 000 (FWHM at *m/z* = 200). Model fitting and other plots were made with GraphPad Prism 9.

## Results and Discussion

### Characterization of IR-MALDESI-MS for high-throughput protein analysis

To establish the detectable protein molecular weight range with our IR-MALDESI-MS system, we analyzed protein standards ranging from 12 kDa to 150 kDa (cytochrome C, carbonic anhydrase, recombinant human serum albumin/rHSA and NISTmAb). The protein standards were either dissolved in water (cytochrome C and carbonic anhydrase), diluted in water from buffered solution (rHSA) or buffer exchanged to 100 mM ammonium acetate (NISTmAb). These mass spectra were collected using regular handshake mode (described previously^13^, see below) and an injection time of 100 ms. Raw mass-to-charge spectra are shown in Figure 1 with deconvoluted mass spectra as insets. Multiple glycoforms were observed in the deconvoluted spectrum of NISTmAb (Figure 1d, inset and Figure S1).

**Figure 1.**
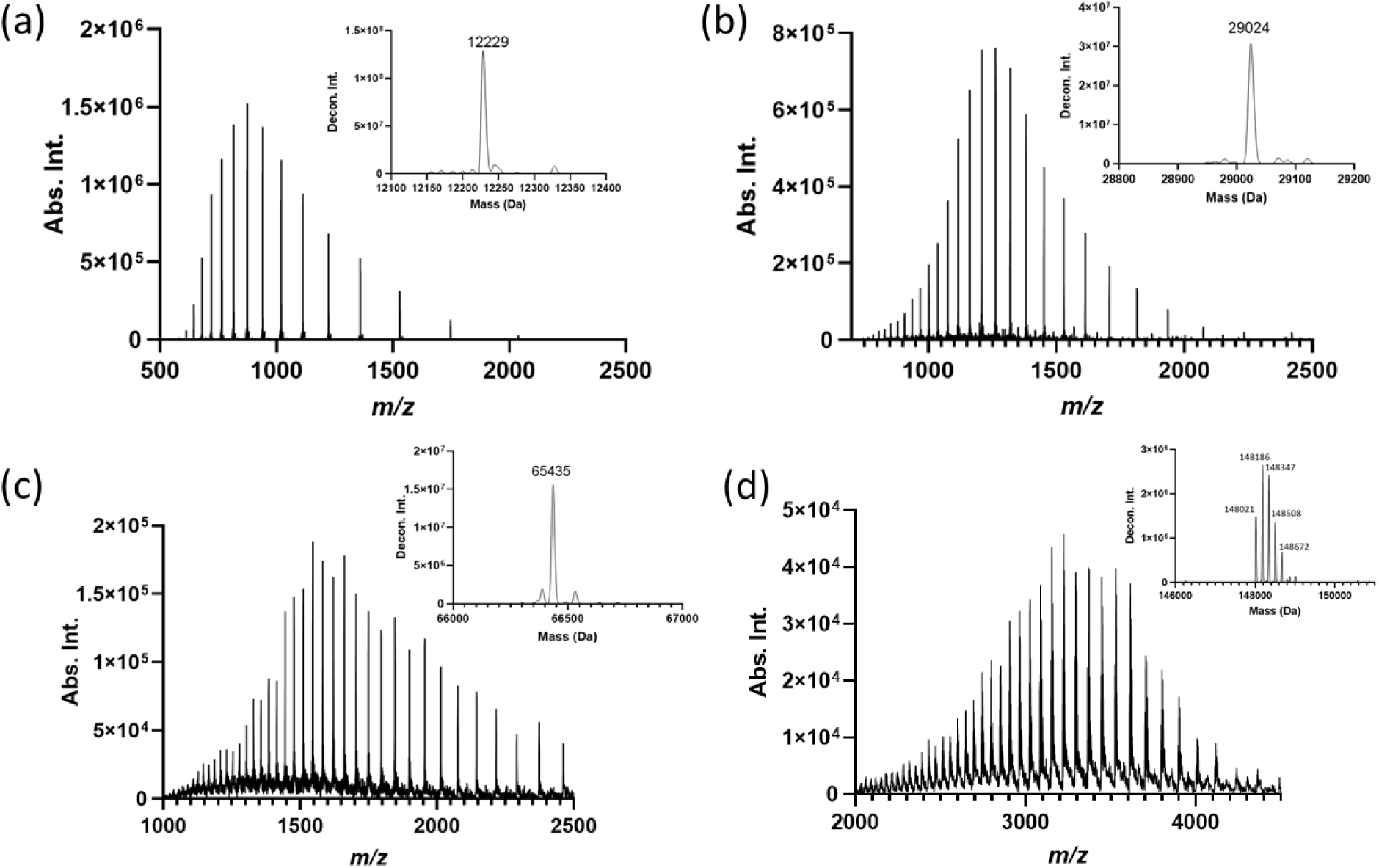
Protein mass spectra of standards collected using IR-MALDESI-MS (30 summed scans). (a) 16 µM cytochrome C; (b) 172 µM carbonic anhydrase; (c) 46 µM recombinant human serum albumin (diluted in water from buffered stock solution); (d) 67 µM NISTmAb (buffer exchanged to 100 mM ammonium acetate). Insets: deconvoluted spectra.

To examine how resolving power (RP) impacted our capability to analyze intact proteins, we first compared the relationship between RP and analysis speed using different scan modes. Injection time was fixed at 20 ms and the sample was a 384-well plate with alternating columns of 5 µM carbonic anhydrase and water. In continuous scan mode, the time to run a full 384 well-plate increased with RP. The fastest speed was achieved with a RP of 7, 500 at 17.5 s per plate (i.e., 22 Hz). Details of this mode have been discussed recently using cytochrome C as the test protein.^15^ Importantly, carryover was not observed, suggesting that the previous sample plume is no longer present when the next sample is ablated. With a larger protein such as carbonic anhydrase, carryover of greater than 2% was periodically observed (5 wells) under these conditions (Figure S2).

Continuous scan mode is inherently limited to acquiring one spectrum per well (or per sample), which limits its usefulness to analyzing only high concentration (∼1 mg/mL) protein samples. For most applications, where salt and buffer are present, or larger proteins are analyzed, we found that spectral summing or averaging is crucial to obtain good signal-to-noise for a wide range of charge states to yield useful deconvoluted data. To reduce post-acquisition data processing complexity that can result from summing scans, we employed a new triggering mode, that we have termed “microscan mode”. The triggering event in microscan mode is similar to regular handshake mode (see Figure S3 for details), except multiple scans (2 – 10 microscans) can be collected and averaged on the data acquisition computer to generate a single spectrum per triggering event. When 5 spectra were collected for each well, microscan mode improved the speed ∼5x compared to regular handshake mode (Figure 2a). Decreasing the RP from 120, 000 to 30, 000 led to faster scan rates (1.5 Hz or 251 seconds per 384-well plate) but no further improvement was observed at lower RP settings. RP did not have any effect on sample rates achieved in regular handshake mode because an uncontrollable ∼ 600 ms timing delay occurs at the completion of a single scan or a set of microscans^13^. Even at 10 microscans, we found that with an injection time of 20 ms, the sampling rate was not affected by the number of microscans collected per sample (Figure S4). Better quality spectra, increased signal-to-noise ratio (Figure S5) and lower coefficient of variation (Figure S4), were obtained as the number of microscans was increased. This suggests that spectra quality may improve further if the number of microscans could be increased beyond 10 microscans, the current instrument limit. For most assays, we selected 5 microscans to obtain good data quality while preventing the laser from overheating.

**Figure 2.**
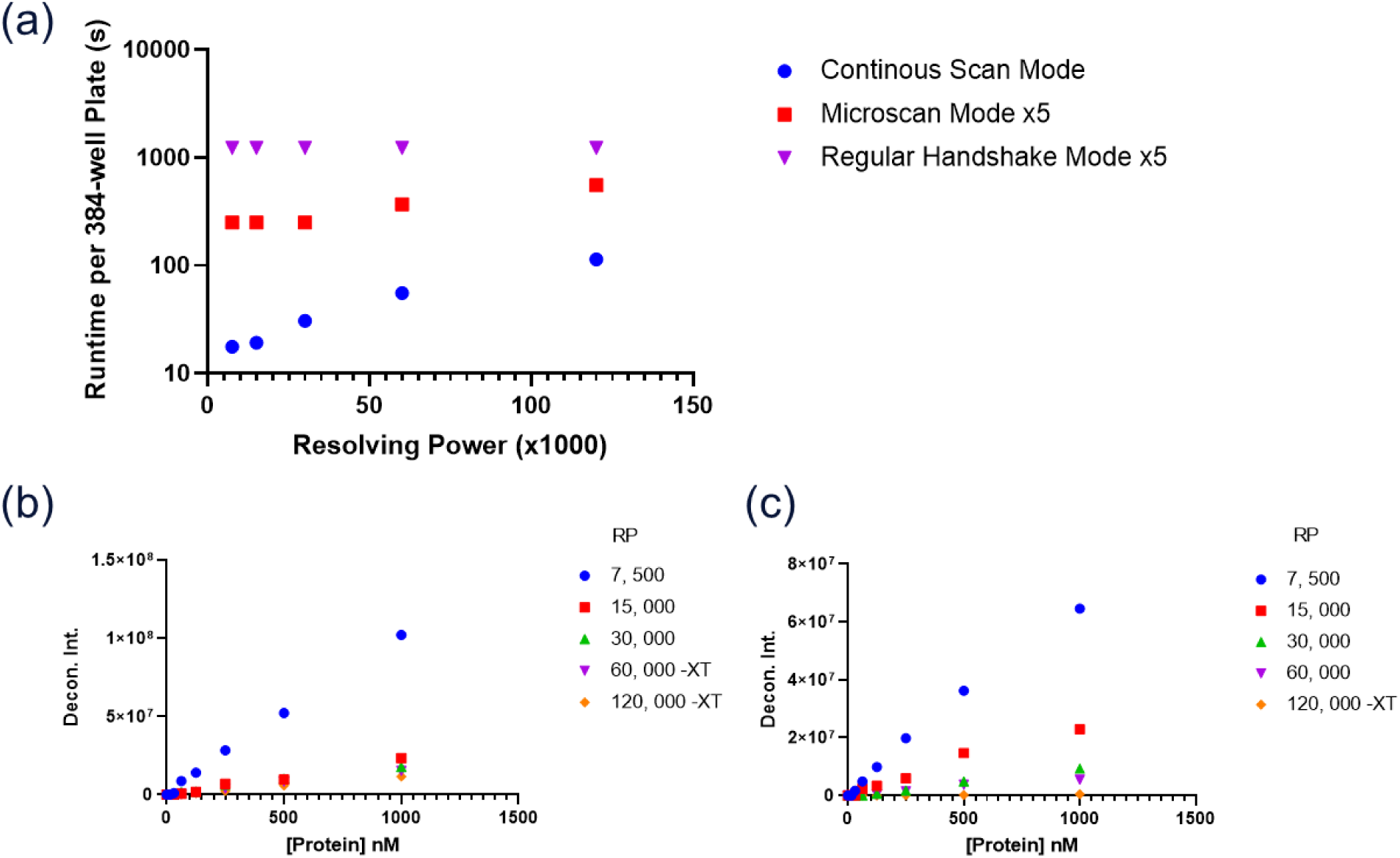
Assesment of the A) acquisition speed and B) sensitivity of cytochrome C and C) sensitiviy of carbonic anhydrase at variours resolution power settings for protein anaalysis by IR-MALDESI. “XT”after RP indicates spectra were deconvoluted using Xtract algorithm instead of ReSpect.

Cytochrome C and carbonic anhydrase standards were prepared at concentrations ranging from ∼15 nM to 1 µM to determine detection limits using microscan mode with 5 microscans per spectrum. The best detection limit (defined as lowest concentration that can be deconvolved with BioPharma Finder) was ∼30 nM for cytochrome C and carbonic anhydrase at a RP of 7, 500 (FWHM) at *m/z* = 200. Mass spectra and deconvoluted spectra of cytochrome C and carbonic anhydrase at 30 nM are shown in Figure S6. This detection limit was achieved with cytochrome C at all tested RPs except for RP of 15, 000. Carbonic anhydrase was only observed at 7, 500 RP. This trend of lower deconvoluted signal was observed for other larger proteins we investigated as well (Figure S7). The true detection limit for IR-MALDESI-MS may be lower, but protein adsorption to the plate walls is significant at nM protein concentrations. We have seen this phenomenon affect some of our enzymatic assays when enzyme concentrations are at low nM levels without the addition of bovine serum albumin or surfactants (data not shown). The addition of these substances is detrimental to analysis of protein analytes at low concentrations. Therefore, well plates with lower protein binding properties would be highly beneficial for IR-MALDI-MS as it is for other protein / peptide LC/MS-centric workflows.

Previously, it was established that the IR-MALDESI source is compatible with complex matrices for small molecule analysis.^13,30^ In this study, we tested signal suppression effects on cytochrome C and carbonic anhydrase (both at 11 µM) from several common buffer components used for storage of target proteins. Three-fold serial titrations of matrices were performed individually with NaCl, MgCl_2_, Tris (Ph = 7.5) and HEPES (pH = 8) starting at 900 mM, Tween-20 at 0.9% and glycerol at 9%. Maximum titrated concentrations of matrices, which could still produce deconvoluted protein mass spectra, were defined as tolerable limits.

The results are summarized in Table S1. We observed concentration-response effects for NaCl, MgCl_2_, Tris and HEPES (Figure S8a, b). These salts and buffers were well tolerated at 30 – 100 mM levels. Tween-20 was not well tolerated and should be avoided or removed during sample preparation. Glycerol had no significant ion suppression effect within the tested range (Figure S8c, d). The buffer tolerance assessments (see above) were performed using 384-well plates with 5 spectra collected per well and then averaged for deconvolution (regular handshake mode, 0.3 Hz). These experiments have served as a guide to assess when a recombinant target protein can be analyzed directly from the storage buffer system for real-time kinetic or compound binding experiments or if dilution with water or exchange into an ammonium acetate buffer is necessary. Fortunately, we have typically obtained satisfactory data when analyzing in-house project target proteins directly from the material as provided or after a 2-3x dilution with water.

### High Throughput Sample Cleanup Coupled with IR-MALDESI-MS

For assays with salt/buffer concentrations that are incompatible with IR-MALDESI or that require low protein concentrations, a simple multiplexed sample cleanup can be deployed. We investigated two multiplexed plate-based strategies and one that utilized magnetic beads to prepare samples for investigation by high throughput protein IR-MALDESI-MS.

Lab-in-a-Plate™ (slit plate) is a commercially available desalting plate-based version of the Top-Tip™ desalting device used for individual samples. Micrometer sized slits, cut at the bottom of wells, are filled with particles coated with chromatographic stationary phase particles. We performed a series of experiments with a C4 slit plate to determine how well 23 µM La protein spiked in PBS (1X) could be desalted using 1 or 3 equilibration and wash cycles. Different equilibration and wash cycles were compared. Samples were eluted to a PCR plate and IR-MALDESI-MS was performed directly from the PCR plate. A significant difference between the mass spectra could be observed before and after being desalted (Figure 3a and 3b). Using the signal-to-noise ratio of the 16^+^ peak (*m/z* ∼797) to assess recovery, it was clear that one equilibration and wash cycle was sufficient to achieve a good signal (Figure S9) even though the vendor recommends three cycles. The increased tolerance to salt provided by IR-MALDESI compared to ESI or MALDI may have made this reduction in sample processing possible. Each equilibrium and wash cycle required a separate centrifugation step, reducing the number of cycles decreased the completion of the process from 30 min to 10 min.

**Figure 3.**
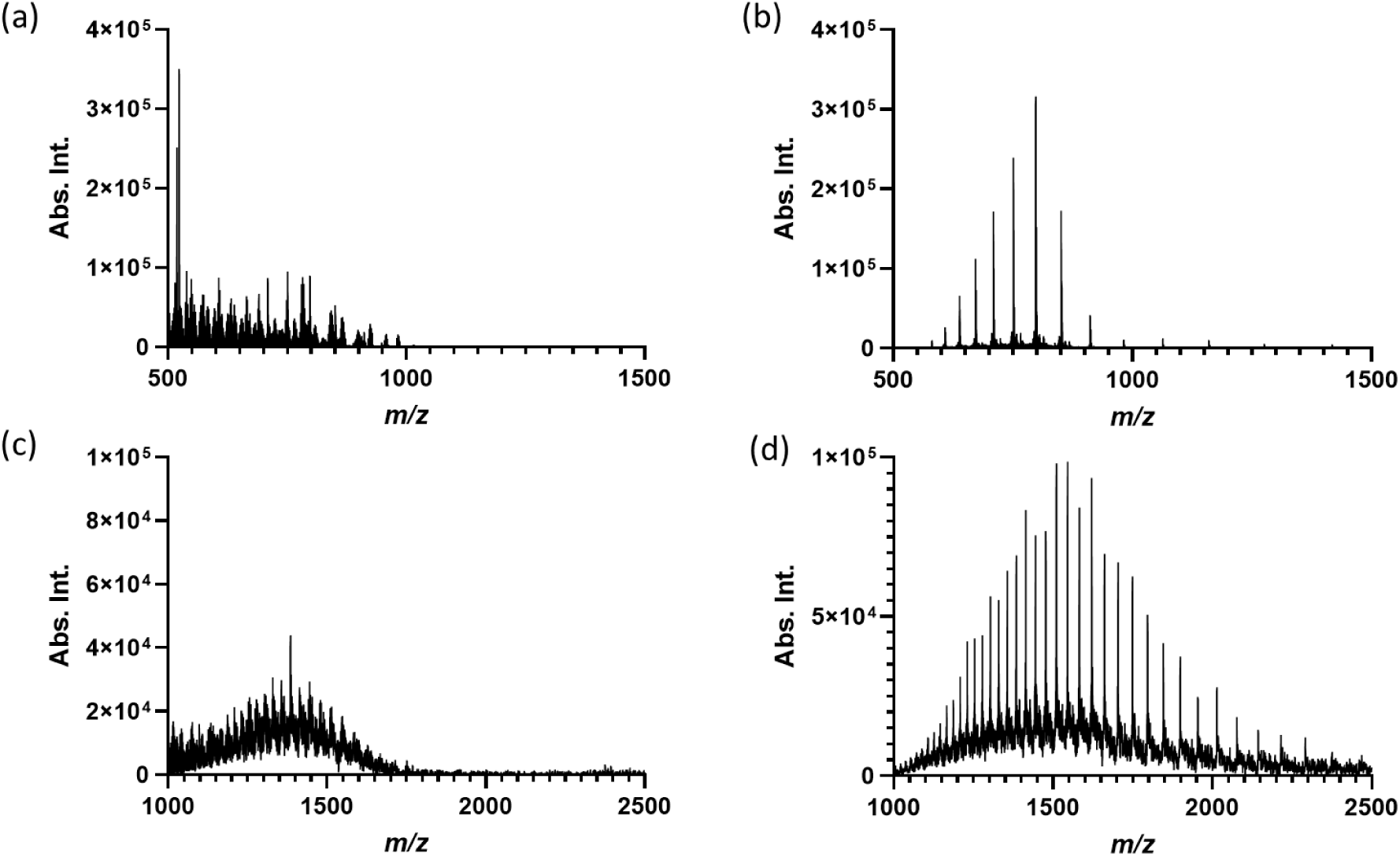
High throughput desalting coupled with IR-MALDESI-MS. Before (a) and after (b) slit plate desalting of La protein in PBS 1X; before (c) and after (d) C-18 magnetic beads desalting of rHSA in PBS 1X.

Magnetic beads are another fast and flexible plate-based sample cleanup method that can be performed within 10 min. The beads can be coated with varied materials such as C18, Ni-NTA, or protein A to either desalt / isolate and concentrate a protein of interest. After protein binding, the beads can be collected with a magnet and washes can be performed to remove unwanted species. The sample cleanup can be automated to work with standard liquid handling equipment. A proof-of-concept experiment using C18 coated magnetic beads (BcMag™ C18) on a Blue®Washer (BlueCatBio) was conducted with 1.5 µM rHSA spiked into PBS (1X) to simulate a protein sample requiring desalting. After this procedure, a discernible protein envelope could be deconvoluted when compared to sampling protein directly from PBS buffer (Figure 3c and 3d). Compared to the Lab-in-a-Plate™ methodology, magnetic bead cleanup with the Blue®Washer does not require any separate centrifugation or dispensing steps and the device can be programmed to evacuate and dispense liquid.

Even with off-line desalting process added to the workflow, IR-MALDESI still requires significantly less time (ca. 14.2 min, 10 min for desalting and 4.2 min when collecting data at a 1.5 Hz sample rate) compared to SampleStream (96 min, 0.04 Hz sample rate)^7^ or the RapidFire system (128 min, 0.05 Hz sample rate)^6^ to analyze 384 well-plates of samples.

To demonstrate the flexibility of using magbeads for sample clean-up, Protein A magnetic beads were used to capture NISTmAb spiked into PBS (Figure S10) to mimic how to monitor for the yield and quality of a therapeutic antibody during a fermentation process^31^.

### Kinetics Measurements: NISTmAb digestion, BTK Phosphorylation and Compound Binding Kinetics

The fast analysis speed of IR-MALDESI can be leveraged to analyze large numbers of samples or to measure fewer samples repetitively. With the tolerance to typical protein buffer systems provided by IR-MALDESI, it is feasible to measure the kinetics of protein – small molecule covalent and non-covalent interactions. A series of experiments were conducted to demonstrate interaction types with various experimental timescales.

A simple kinetic measurement experiment was performed to monitor NISTmAb reduction by 5 mM TCEP (tris(2-carboxyethyl) phosphine) from the intact form to its light and heavy chain components. Measurements were taken repetitively from the same well using regular handshake mode with an injection time of 100 ms (five scans per sample, 0.3 Hz). The initial data point was taken ∼20 s after TCEP was added to NISTmAb. A shift in heavy chain/light chain ratio was observed. Light chain was detected at the first time point while heavy chain was still not identified (Figure 4a). Peaks corresponding to two linked heavy chains could be found at early time points and were subsequently converted to the single heavy chain species (Figure 4a-c). The transition was completed in about 10 min and the heavy chain/light chain ratio remained stable from that point (Figure 4d) with heavy chain glycoforms mass-resolved after deconvolution (Figure S11).

**Figure 4.**
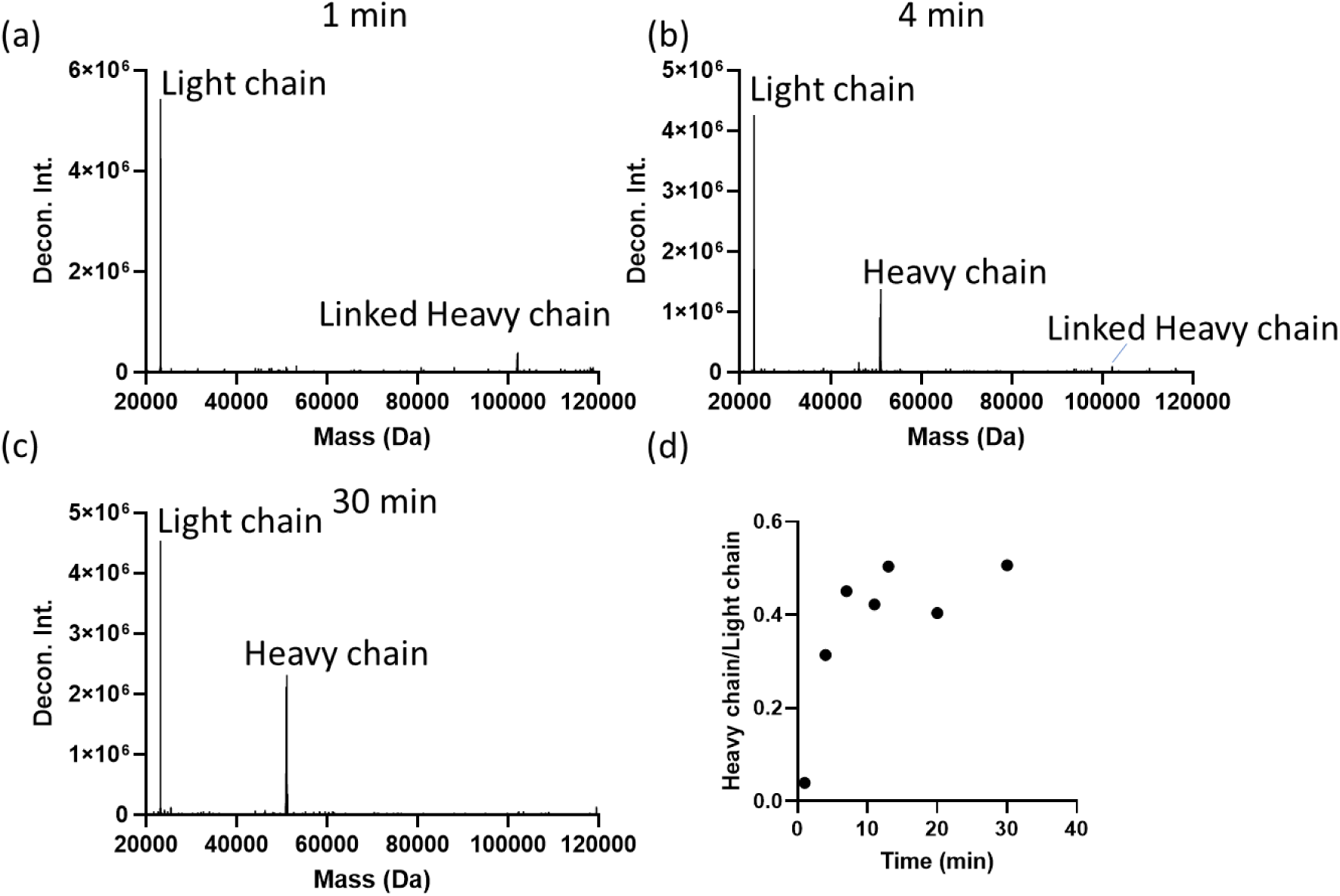
(a)-(c) Deconvoluted spectra at 1 min, 4 min and 30 min after adding TCEP to NISTmAb; (d) Heavy chain/light chain ratio after adding TCEP.

To further illustrate the near-real time sampling capability of IR_MALDESI, the autophosphorylation rate of recombinant purified Bruton’s tyrosine kinase (BTK) kinase domain was monitored. BTK was measured directly in 5 mM HEPES with IR-MALDESI-MS. Using the provided sample, unphosphorylated BTK was observed with a deconvoluted mass of 33, 505 Da (Figure 5a). The detection limit of BTK protein was established by a three-fold serial titration experiment, which was ∼400 nM in 5 mM HEPES buffer (Figure 5b). There is one phosphorylation site in the BTK kinase domain (Y551),^32^ and we observed only one addition of 80 Da (33, 585 Da) during the entire time course experiment (Figure 5c). The progress of autophosphorylation was represented by percent mono-phosphorylated (Figure 5d), which was calculated as the ratio of deconvoluted intensity of mono-phosphorylated (reacted) BTK over the sum of unreacted and reacted BTK protein. We did not observe any phosphorylated BTK until the reaction reached 10% conversion. All data were collected by repetitively sampling from just three samples. Autophosphorylation was completed in ∼3 hours under these experimental conditions.

**Figure 5.**
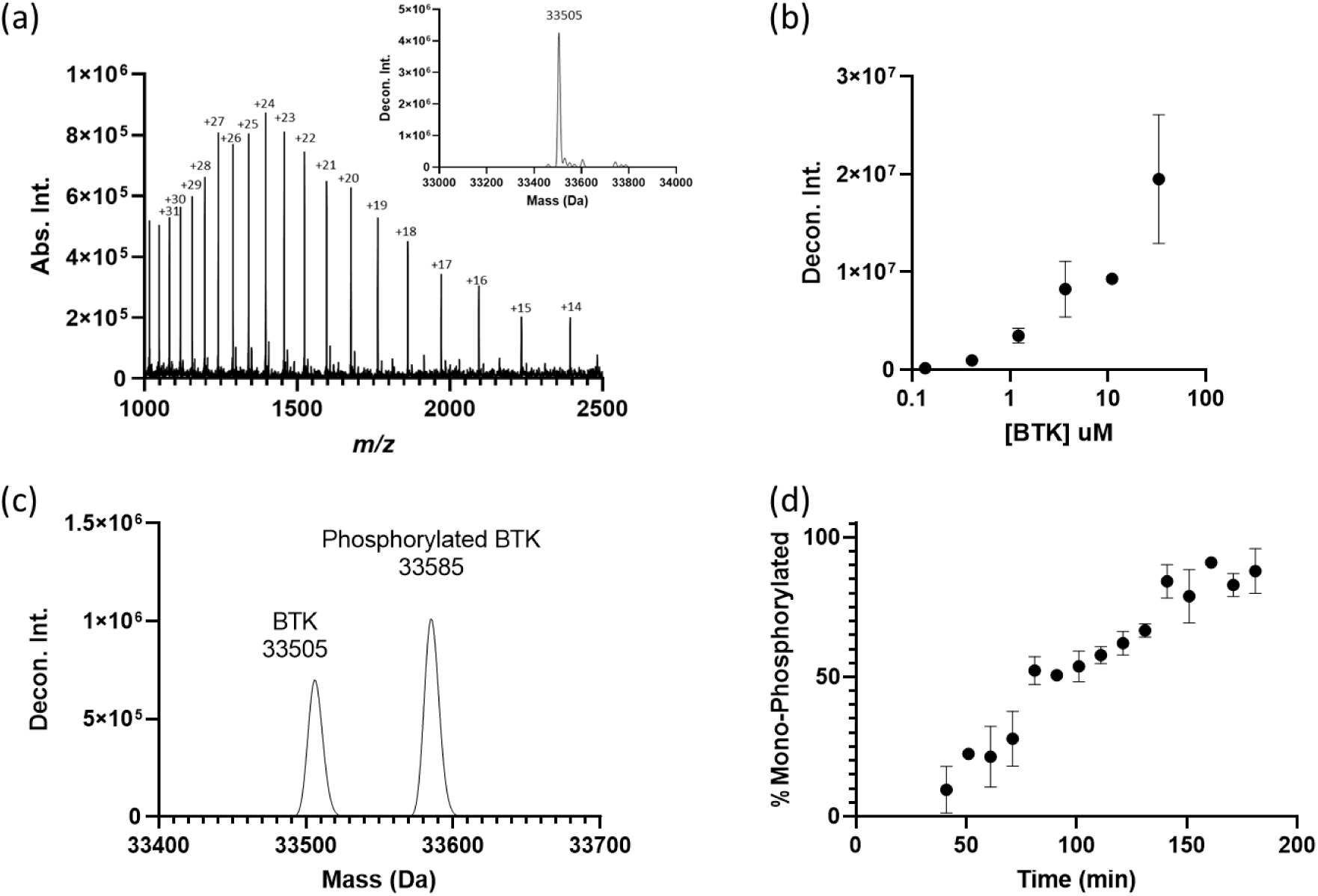
(a) Raw mass spectrum and deconvoluted spectrum (inset) of BTK protein; (b) standard curve of BTK protein in 5 mM HEPES buffer using deconvoluted intensities; (c) deconvoluted spectrum of BTK autophosphorylation in progress; (d) time course of BTK autophosphorylation measured by IR-MALDESI-MS in kinetic mode.

There is renewed interest in covalent inhibitors as potent and effective agents, despite having been deprioritized in drug development in the past due to safety concerns that now can be mitigated through careful drug compound design^33^. Several covalent inhibitors and non-covalent inhibitors for BTK have been discovered and are approved or in advanced clinical stages^34^. We selected four compounds: acalabrutinib and zanubrutinib (irreversible covalent), rilzabrutinib (reversible covalent), and vecabrutinib (non-covalent) to measure their binding behavior to BTK using IR-MALDESI-MS. Figure S12 shows deconvoluted spectra after incubating BTK with compounds for 10 min at 1:1 molar ratio. There were no peaks corresponding to compound addition for vecabrutinib (Figure S12a) since the base peak mass was consistent with unmodified intact protein (33505 Da). This indicated that under these experimental conditions non-covalent complexes were not preserved. However, for BTK binding with acalabrutinib (MW 465 Da), zanubrutinib (MW 471 Da) and rilzabrutinib (MW 665 Da), the base peak mass increased to 33970 Da, 33976 Da, and 34370 Da, respectively (Figure S12b-d), which was consistent with each compound covalently binding with BTK.

Lowering the pH of the protein-compound mixture with an equal volume of 1% formic acid caused dissociation of the reversible covalent complex (BTK-rilzabrutinib) that could be measured in real-time by IR-MALDESI-MS. We observed the compound adduct peak gradually converted to the intact protein peak represented by the reduction of percent bound compound (Figure 6a, calculated as deconvoluted intensity of bound protein over the sum of bound and unbound protein). As expected, the complex formed between BTK, and the irreversible covalent compounds remained intact over a similar period (Figure 6b).

**Figure 6.**
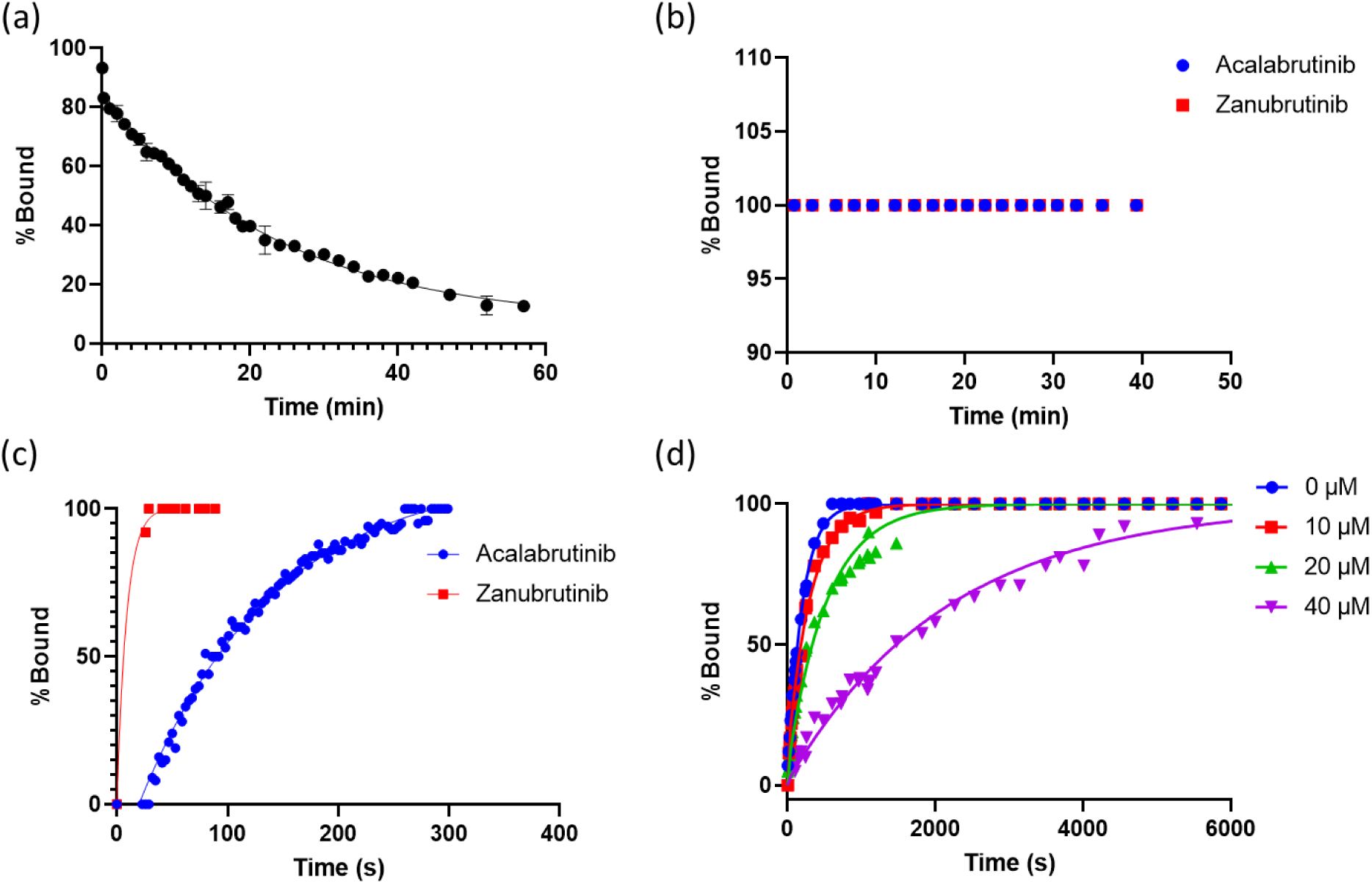
(a) Denaturing of rilzabrutinib and BTK complex via lowering pH; (b) lowering pH did not reverse acalabrutinib/zanubrutinib binding to BTK; (c) monitoring fast covalent binding of acalabrutinib and zanubrutinib to BTK; (d) competitive binding experiment by adding vecabrutinib and acalabrutinib simultaneously to BTK protein, concentrations of vecabrutinib was a variable and reflected as different curves in this figure, K_D, vecabrutinib_ = 30 nM.

Although not demonstrated here, we are assessing if IR-MALDESI-MS can analyze less stable reversible covalent complexes (e.g., fast reversible) that are undetectable by LC/MS. Protein denaturation, due to prolonged exposure to formic acid and acetonitrile in the mobile phase when on the LC column, may destabilize the binding site and lead to dissociation of the reversible protein-compound bond resulting in detection of only unmodified protein by the mass spectrometer. For IR-MALDESI-MS, dissociation may be minimized or eliminated since the reversible covalent complex is exposed to denaturing solvents in the ESI spray on a millisecond time scale.^35^

It is possible to monitor fast covalent binding events by using microscan mode, which is demonstrated in Figure 6c. Three spectra were collected for each data point with an injection time of 100 ms, leading to a time resolution of 3s. The reaction was initiated manually in a non-reading position, therefore there was 20-30 s overhead time before the first data point was taken. This overhead time can be eliminated by devices that dispense reagent automatically at the IR-MALDESI reading position. We were unable to capture an association curve for zanubrutinib because the binding ratio reached 100% within 30 s. However, a complete association curve for acalabrutinib can be obtained and the binding ratio reached 100% in ca. 5 min.

Due to the irreversible covalent binding of acalabrutinib to BTK, it can serve as a probe to provide insights to kinetics of other non-covalent and reversible covalent binders. In this experiment, vecabrutinib and acalabrutinib were added to BTK solution simultaneously (Figure 6d). The concentration of acalabrutinib was held at a constant 200 µM whereas the concentration of vecabrutinib was varied from 0 to 40 µM and placed in four separate wells on the plate. By monitoring the formation of the acalabrutinib-BTK complex, vecabrutinib binding kinetics can be determined by the Motulsky-Mahan model^36^. In the absence of vecabrutinib, acalabrutinib binding to BTK reached maximum in less than 10 min under current assay conditions, whereas the process was significantly slowed down in the presence of vecabrutinib. This allows association/dissociation rates to be estimated for fast binding reactions that could not be monitored without competition. The estimated association (K_on_) and dissociation rate constant (K_off_) of vecabrutinib was ∼10^6^ M^-1^s^-1^ and 0.03 s^-1^, respectively. K_D_ can be calculated as K_off_/K_on_, which was ∼30 nM.

## Conclusions

In this work, we demonstrated the use of a high throughput IR-MALDESI-MS system to perform protein analysis. Proteins up to 150 kDa could be detected and we benchmarked sample rate, sensitivity, and matrix effects caused by several common buffer components. By using microscan mode, the run time of a 384-well plate was 4.2 min, which is a sample rate of 1.5 Hz. Samples rates up to 22 Hz can be utilized for higher concentrations (µM level) of pure protein. High throughput desalting strategies, based on multi-well microtiter plates, were applied to clean up samples in a 384 well plate without significantly sacrificing throughput using multiple techniques. Besides analyzing a large quantity number of samples, high analysis speed was utilized to study kinetics, as demonstrated by monitoring TCEP reduction of NISTmAb along with BTK autophosphorylation and compound binding experiments. The most significant bottleneck we are experiencing is producing deconvoluted mass spectra, since it is done on a sample-per-sample basis rather than on a plate-per-plate basis. Besides applications discussed in this work, we have also explored additional applications to support drug discovery efforts, such as covalent fragment screening and compound induced target protein modifications (binding, oxidation, or aggregation), and additional triggering schemes to increase throughput. We believe IR-MALDESI-MS is a versatile high throughput platform that can be used widely for protein analysis applications.

## Acknowledgements

The authors acknowledge Laura J. Miesbauer, Robert W. Johnson and Kevan Knizner for their critical insights. The work was enabled by the AbbVie Postdoc Program. All authors are employees of AbbVie. The design, study conduct, and financial support for this research were provided by AbbVie. AbbVie participated in the interpretation of data, review, and approval of the publication.

## Notes

### Summary of Updates

Restructured to include new data and removed some data that did not fit in

