## Supporting Information for "High Throughput Intact Protein Analysis for Drug Discovery Using Infrared Matrix-Assisted Laser Desorption Electrospray Ionization Mass Spectrometry"

This file includes 15 pages: 12 figures (pages S3-S15) and supplementary text (page S13)

### Characterization of IR-MALDESI-MS for High-throughput Protein Analysis

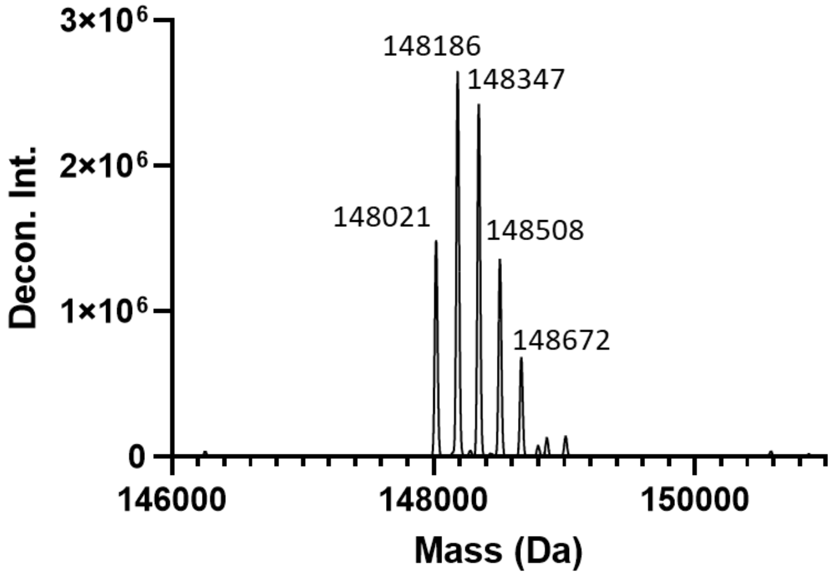

*Figure S 1. Deconvoluted spectrum of intact NISTmAb shows different glycoforms.*

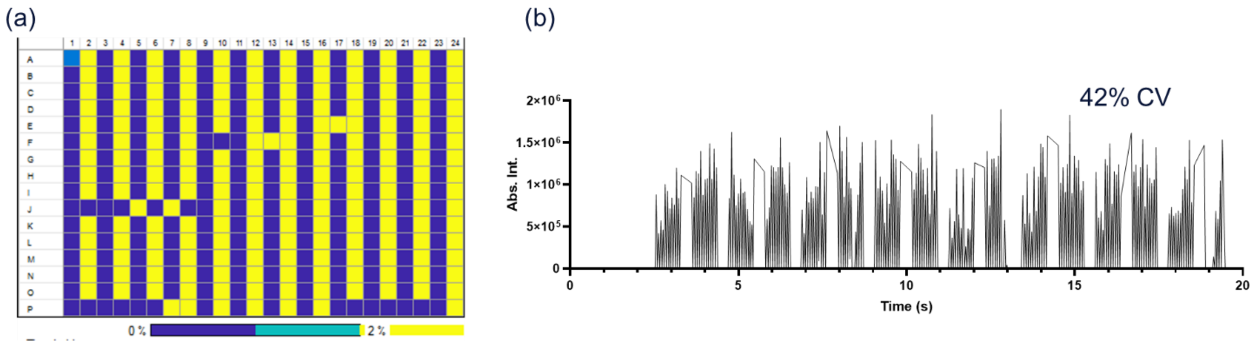

Figure S 2. Analysis of a 384-well plate with alternating columns of 5 µM carbonic anhydrase and water. (1) Heatmap of m/z 1531.87 (z = 19) with 200 ppm mass tolerance. Yellow indicates detected at >2% of maximum intensity on the plate. (2) Extracted ion chromatogram (EIC) of m/z 1531.87 with 200 ppm mass tolerance, plate reading time was 17.5 s.

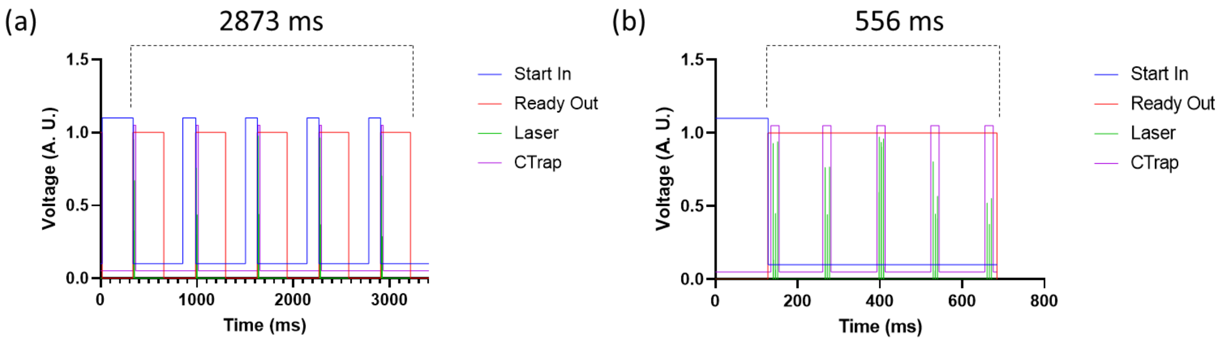

Figure S 3. Triggering events during IR-MALDESI acquisition under (a) regular handshake mode and (b) microscan mode. MS resolving power: 60, 0000 (FWHM @ m/z 200), injection time: 20 ms. Start in: signal sent to MS to initiate a scan; ready out: signal returned from MS indicating MS is acquiring; Laser: laser fires; C Trap: C Trap opens.

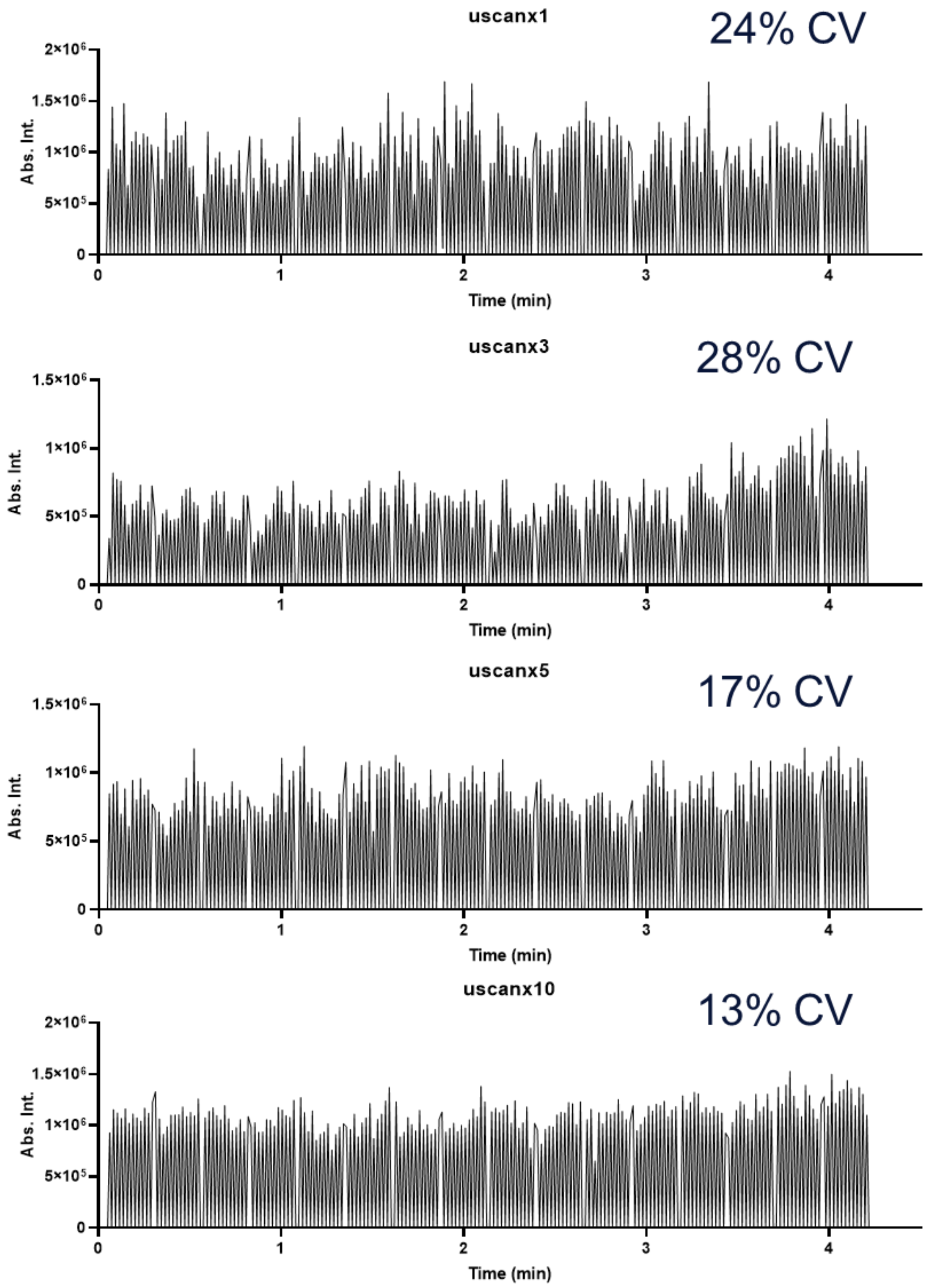

Figure S 4. Increasing the number of microscans does not affect the scan speed but with more microscans %CV decreases. From top to bottom: 1, 3, 5 and 10 microscans per spectrum.

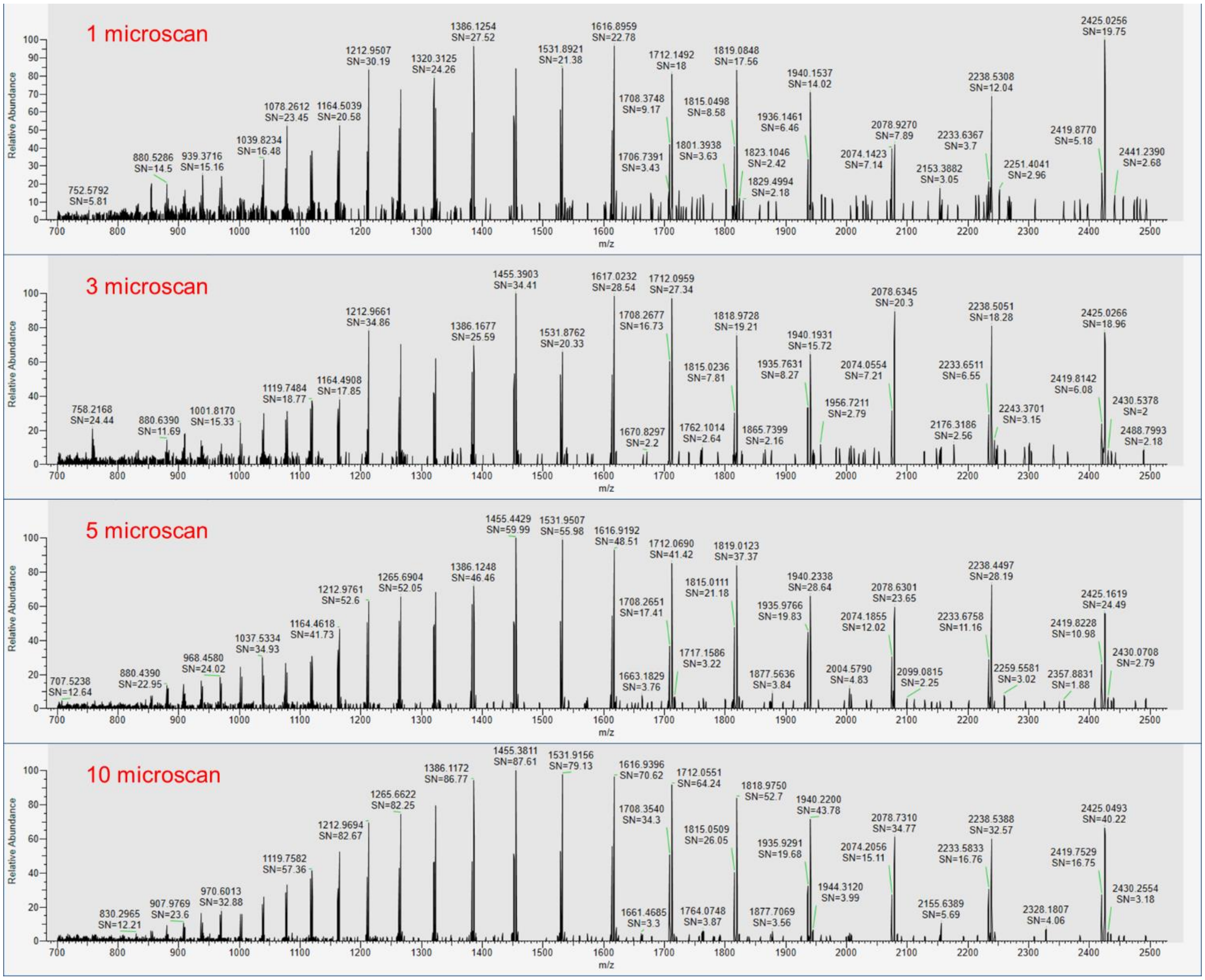

Figure S 5 Increasing the number of microscan yields better spectra. Top to bottom: 1, 3, 5 and 10 microscans per spectrum.

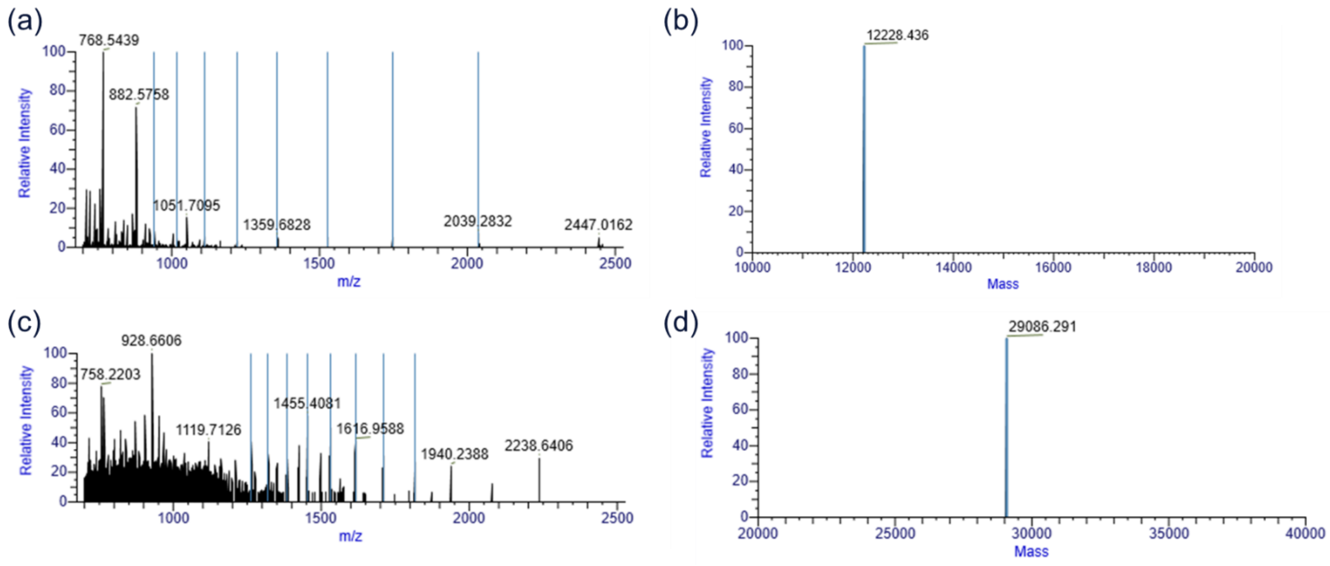

Figure S 6. (a) Raw mass spectrum and (b) deconvoluted spectrum of 30 nM cytochrome C; (c) Raw mass spectrum and (d) deconvoluted spectrum of 30 nM carbonic anhydrase.

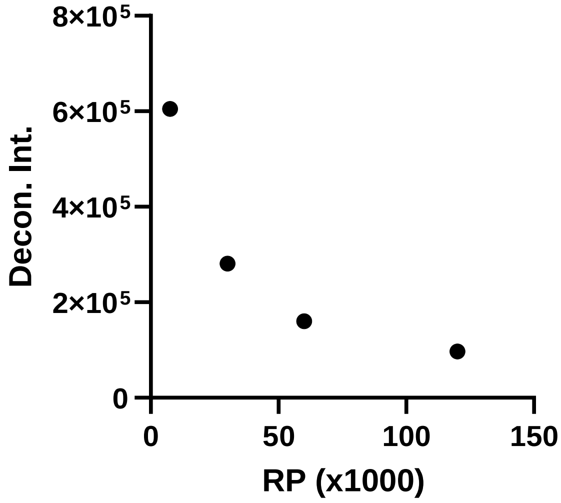

Figure S 7. Deconvoluted intensity of 67 µM NISTmAb decreased with increased resolving power.

Table S1. IR-MALDESI-MS matrices tolerance for Cytochrome C and Carbonic Anhydrase

| Tolerable Limits | Cytochrome C | Carbonic Anhydrase |
|---|---|---|
| NaCl (mM) | 300 | 100 |
| $MgCl_2$ (mM) | 100 | 100 |
| Tris (pH 7.5) (mM) | 300 | 300 |
| HEPES (pH 8) (mM) | 100 | 33 |
| Tween-20 (%) | 0.01% | N/A |
| Glycerol (%) | 9% | 9% |

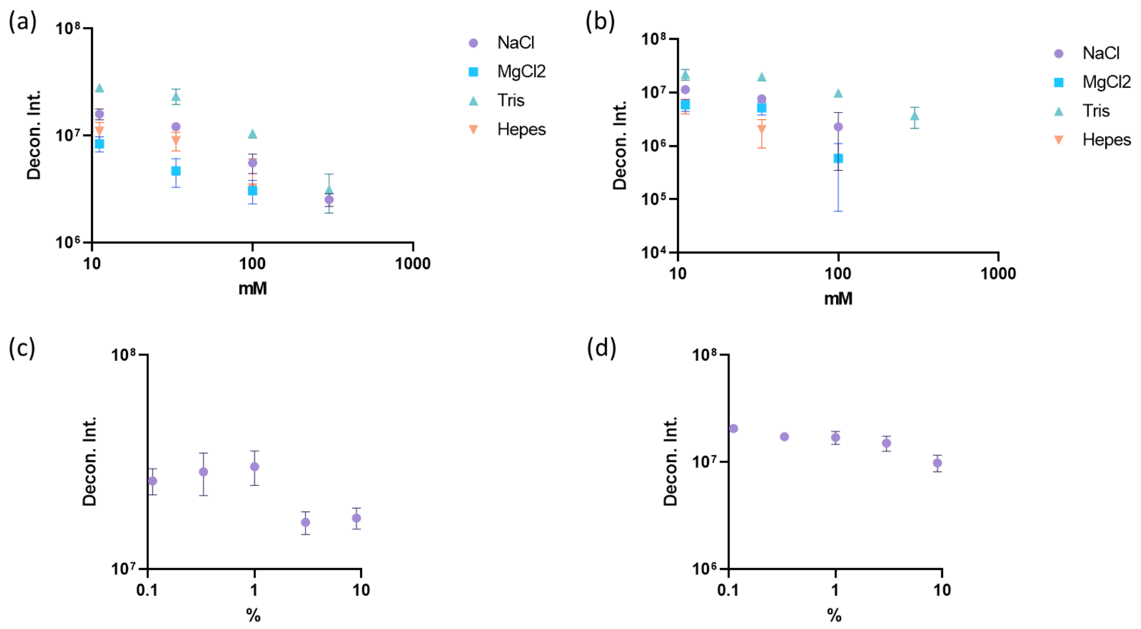

Figure S 8. Effect of salts/buffers on IR-MALDESI-MS detection of (a) cytochrome C and (b) carbonic anhydrase. Effect of glycerol on detection of (c) cytochrome C and (d) carbonic anhydrase.

### High Throughput Sample Cleanup Coupled with IR-MALDESI-MS

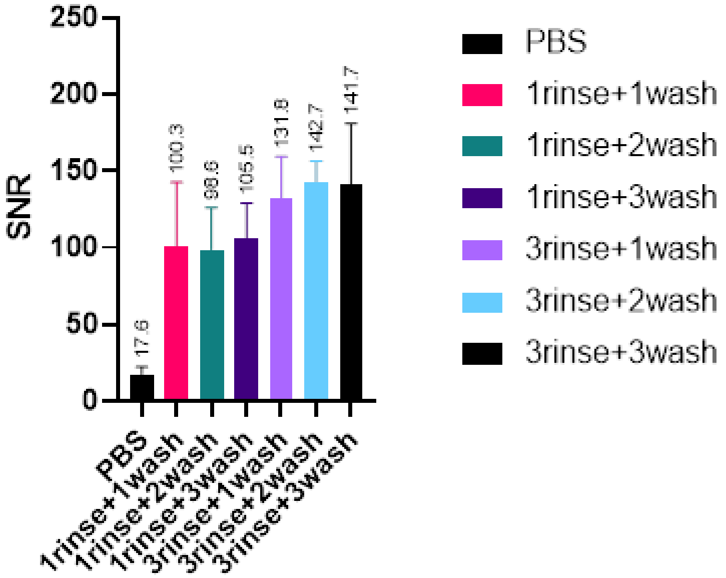

Figure S9. Comparison of different wash protocols using slit plate. SNR of m/z 797 was used to assess signal recovery.

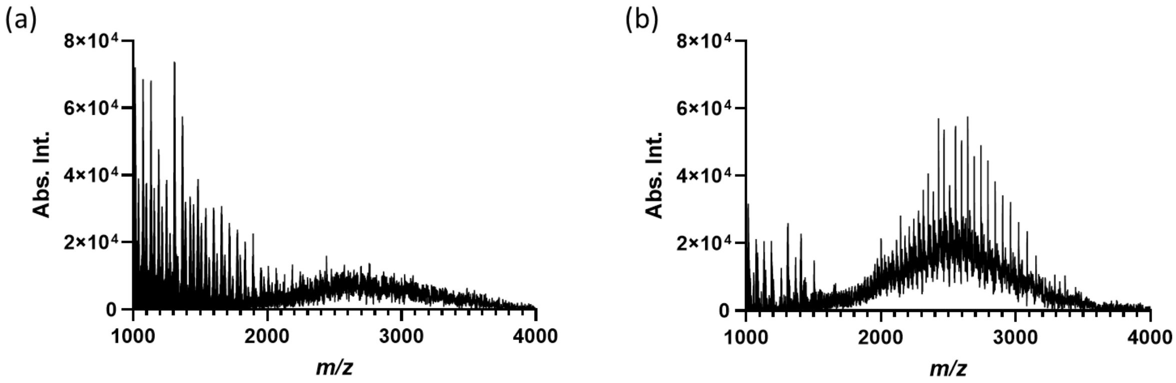

Figure S 10. Protein A magnetic bead cleanup of NISTmAb. (a) 1.44 mg/mL NISTmAb in PBS; (b) same sample after cleaning with Protein A magnetic beads.

### Sample Cleanup with Protein A Magnetic Beads

Sample cleanup with Protein A magnetic beads was performed according to vendor protocol, except 20 mM Tris pH 7.5 was used as wash/binding buffer and 10 mM HCl was used as elution buffer. Briefly, 50 µL of Pierce Protein A magnetic beads were mixed with 150 µL wash buffer, beads were then collected by a magnet and supernatant was removed. After one additional wash, the sample was diluted in wash buffer (100 µL sample + 400 µL wash buffer) and mixed with beads. The mixture was then incubated at room temperature with mixing for 1 h. After incubation, beads were washed three times with wash buffer then 100 µL of elution buffer was added and incubated for 15 min. The beads were then collected by magnet and eluent was transferred to microtiter plate for IR-MALDESI-MS.

### Kinetics Measurements: NISTmAb digestion, BTK Phosphorylation and Compound Binding Kinetics

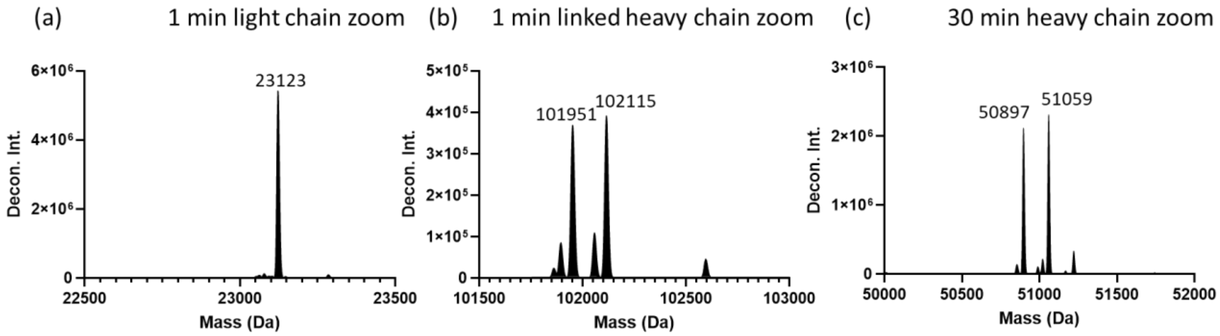

Figure S 11. Zoomed-in view of deconvoluted spectra at different time points after adding TCEP to NISTmAb, glycoforms could be resolved for heavy chains.

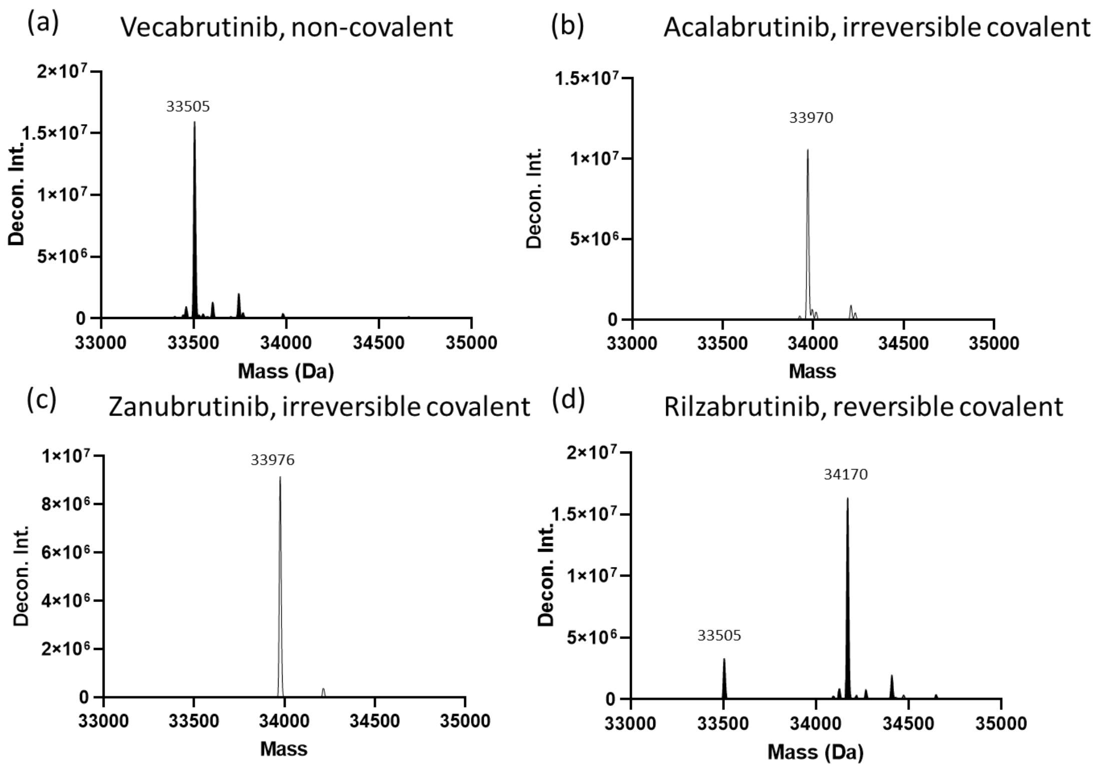

Figure S 12. BTK binding with different compounds as measured by IR-MALDESI-MS. (a) Vecabrutinib; (b) acalabrutinib; (c) zanubrutinib and (d) rilzabrutinib.
